# Biphasic Mechanical Loading Disrupts Cytoskeletal Symmetry in 3D Architected Scaffolds

**DOI:** 10.1101/2025.08.04.668203

**Authors:** Kailin Chen, Alexander Bolanos-Campos, Mistica Lozano Perez, Erin Berlew, Tianbai Wang, Luc Capaldi, Ran Tao, Arnold Mathijssen, Joel D. Boerckel, Ottman A. Tertuliano

## Abstract

Cells in load-bearing tissues experience both solid deformation and interstitial fluid flow during physiological loading, but the mechanisms by which they integrate these biphasic mechanical signals remain poorly understood. Here, we develop a porous, nanoarchitected 3D scaffold that allows simultaneous delivery and control of matrix strain and fluid shear stress. We validated the platform through fatigue loading experiments and simulations of fluid–structure interactions. In static culture, osteoblast-like cells adopted shapes, cytoskeletal architectures, and focal adhesion patterns templated by scaffold geometry. Under cyclic compression, the combined influence of matrix deformation and induced fluid flow disrupted this alignment, producing disordered actin structures and reduced focal adhesion eccentricity. These changes emerged even under low-frequency loading, within the drained poroelastic regime, indicating a high sensitivity of cytoskeletal organization to fluid-solid coupling. Our findings establish a tractable and tunable platform to investigate how cells sense and respond to dynamic biphasic mechanical environments in 3D.

**Significance Statement:** Cells in tissues such as bone experience mechanical inputs from both matrix deformation and interstitial fluid flow. However, existing in vitro systems often isolate one type of input or lack the ability to control both independently. We engineered a nanoarchitected 3D scaffold that delivers tunable biphasic mechanical inputs by combining structural compression and fluid flow. Without external loads, cells align their cytoskeleton and focal adhesions to the scaffold geometry. When subjected to dynamic loading, they transition to disordered morphologies and less mature focal adhesions, suggesting a transition to migratory states. These results highlight the sensitivity of cells to even subtle biphasic cues and provide a new platform to study how cells integrate multiple mechanical signals in 3D environments.

## Introduction

Cyclic deformation of tissues causes extracellular matrix deformation, interstitial fluid flow and dynamic mechanical stimuli of the residing cells. In bone, cartilage, and other load-bearing tissues, these mechanical inputs are inherently biphasic—driven by the poroelastic coupling between a viscoelastic solid scaffold and a permeating viscous fluid^1–3^. These biphasic cues regulate diverse cellular behaviors like cytoskeletal remodeling^4^, focal adhesion formation^5^, and cell migration^6^. However, it remains unclear how cells interpret and respond to these coupled solid and fluid mechanical signals because control of such complex mechanical environments in vitro has been experimentally inaccessible.

While 2D culture systems have provided key insights into mechanotransduction^7,8^, they fail to recapitulate the geometric confinement and fluid resistance encountered by cells in vivo. 3D matrices and organoid models capture some aspects of spatial confinement and dynamic remodeling^6,9,10^, but typically lack tunable control over the degree of solid–fluid coupling. Most systems are not designed to isolate or quantify how matrix strain and fluid shear stress over different time scales result in a single biphasic mechanical stimulus to cells^1,2^.

Cytoskeletal architecture and focal adhesion dynamics are key determinants of how cells sense, integrate, and respond to these stimuli^4,11^. These structures define cell shape and also drive essential processes such as migration, differentiation, and ECM remodeling^7,11^. In 3D environments, cytoskeletal symmetry and organization can be influenced by geometric confinement^12,13^, matrix compliance^8,14^, and fluid flow^6^, and are further modulated by dynamic loading^15,16^. However, few systems offer fine-grained control over scaffold architecture while allowing mechanical inputs to be modulated across physiologically relevant time scales.

To show how cells respond to coupled solid and fluid cues, experimental platforms require control over both geometry and loading dynamics. Architected materials with well-defined geometries and porosity offer an opportunity to investigate how cells align, adhere, and reorganize in response to changes in biphasic mechanical stress^17–19^. For example, studies in 2D fluid shear models suggest that morphological responses to shear gradients depend on adhesion geometry and cytoskeletal tension^20^. These systems usually focus on loading regimes the frequency^21^ or amplitude^22^ of deformation results in only fluid flow^23^ or matrix strain^16^ as the mechanical stimulus, physiologically reminiscent of single overload type injury. However, an important but underexplored regime is the fatigue-relevant, low-frequency poroelastic response of the extracellular matrix, where fluid redistribution remains near equilibrium but may still induce changes in cell response^1–3^. The sensitivity of cells to small perturbations from this drained regime is largely unknown.

Here, we developed a 3D nanoarchitected platform to interrogate cytoskeletal organization under tunable biphasic mechanical loading. We fabricated highly porous scaffolds composed of tetrakaidecahedral unit cells that can each be occupied by an individual cell. By cyclically compressing the scaffolds at frequencies ranging from 0.1–3Hz, we modulate the relative contributions of solid deformation and fluid shear stress experienced by the residing osteoblast-like SAOS-2 cells. Through confocal microscopy of cytoskeletal and focal adhesion protiens and finite element simulations of fluid–structure interactions, we demonstrate that increasing fluid-solid coupling, even within the drained poroelastic regime, disrupts cytoskeletal symmetry. Cells transition from well-defined morphologies that reflect scaffold architecture to irregular shapes characterized by disordered actin and reduced vinculin eccentricity, signaling a loss of focal adhesion maturity. Our findings reveal a previously unrecognized sensitivity to low-level biphasic coupling and provide a tractable system to study cell mechanotransduction dynamically tunable biphasic 3D environments.

## Results

### Solid deformation induces biphasic stresses in 3D structures

We fabricated nanoengineered latticess to serve as a biphasic scaffold for 3D cell culture and mechanical stimulation (Fig. 1A,). Extended fabrication details are in Methods; briefly, using two-photon lithography^24,25^, we first printed polymer scaffolds measuring 125 × 125 × 50 µm^3^ and composed of repeating tetrakaidecahedral unit cells (25 µm wide) with struts approximately 2 µm in thickness. The scaffolds were coated with a 50 nm layer of TiO_2_ by atomic layer deposition (ALD), followed by sacrificial oxygen plasma etching to remove the polymer core, resulting in hollow TiO_2_ shell struts (Fig. S1). These 3D nanoarchitectures were used as in vitro platforms for cell culture under biphasic mechanical conditions.

**Figure 1:**
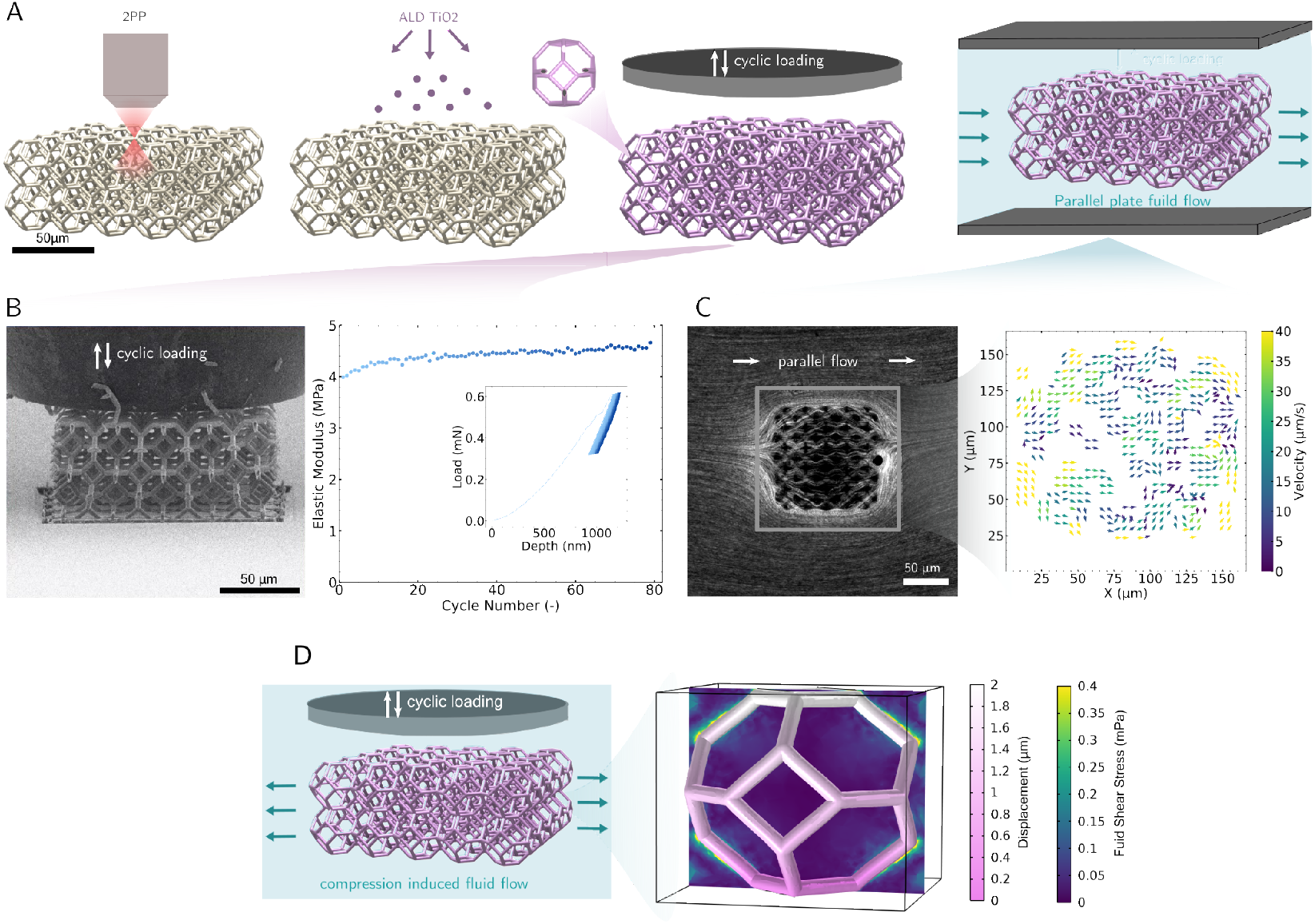
Fabrication and characterization of biphasic nanoarchiteched platform. A) Nanoarchitectures were fabricated using two-photon lithography and coated with TiO_2_ using ALD. Hollow struts were obtained by exposing the polymer with FIB and etching in O_2_ plasma. B) Cyclic compression of nanoarchitectures with hollow struts shows elastic response across 80 cycles. C) Streamlines and velocity field through the nanoarchitecture with parallel flow, measured by µPIV. C) Schematic of cyclic loading on the nanoarchitecture in culture media and the corresponding simulation of fluid shear stress magnitude induced by the solid deformation.

To evaluate the platform’s suitability for biphasic loading, we independently characterized its solid and fluid mechanical behavior. First, we quantified the elastic response of the solid scaffold using cyclic compression tests with an in situ SEM nanoindenter (Fig. 1B, Movie S1). The scaffolds exhibited elastic moduli of 4.5 ± 1.5 MPa and remained structurally stable over 80 cycles without degradation in mechanical response.

Next, we examined fluid transport through the structure by applying laminar flow in a microfluidic channel and visualizing the flow using fluorescent tracer particles (Fig. 1C). Velocity fields were extracted by micro particle image velocimetry (µPIV), revealing flow velocities on the order of 10 µm/s through the lattice. Streamlines followed the scaffold geometry, with localized mixing observed near the nodes. These experimental flow patterns qualitatively matched results from finite element simulations(Fig. S2), though small vortices observed in µPIV were absent in the simulation, likely due to instabilities in the fluid supply. Overall, the agreement suggests that experimentally relevant flow fields and their associated shear stresses can be reasonably captured by simulation.

To characterize the mechanical environment experienced by cells during biphasic loading, we modelled fluid-solid interactions in the scaffold during compression in culture media (Fig. 1D) and developed a complementary 2D squeezing flow framework to estimate fluid shear stress distributions under compressive loading (Fig. S3 - S6). Finite element simulations showed that compression-induced deformation of the solid lattice generates localized shear stresses in the interstitial fluid, particularly near surfaces of the struts. These simulations revealed gradients in shear stress with maxima at strut edges and decreasing toward the center of each pore. The results provide a full-field description of the biphasic stress environment that cells experience within the scaffold and validate this architecture as a platform for probing the coupled effects of matrix deformation and fluid flow in 3D.

### Cytoskeletal organization conforms to nanoarchitecture geometry in the absence of external loading

To understand cellular responses in the absence of external stress, we first examined SAOS-2 cell morphology and cytoskeletal organization on nanoengineered 3D scaffolds under static culture conditions. Cells were seeded at both low and high densities to capture representative single-cell and population-level behavior (Fig. 2A,B).

**Figure 2:**
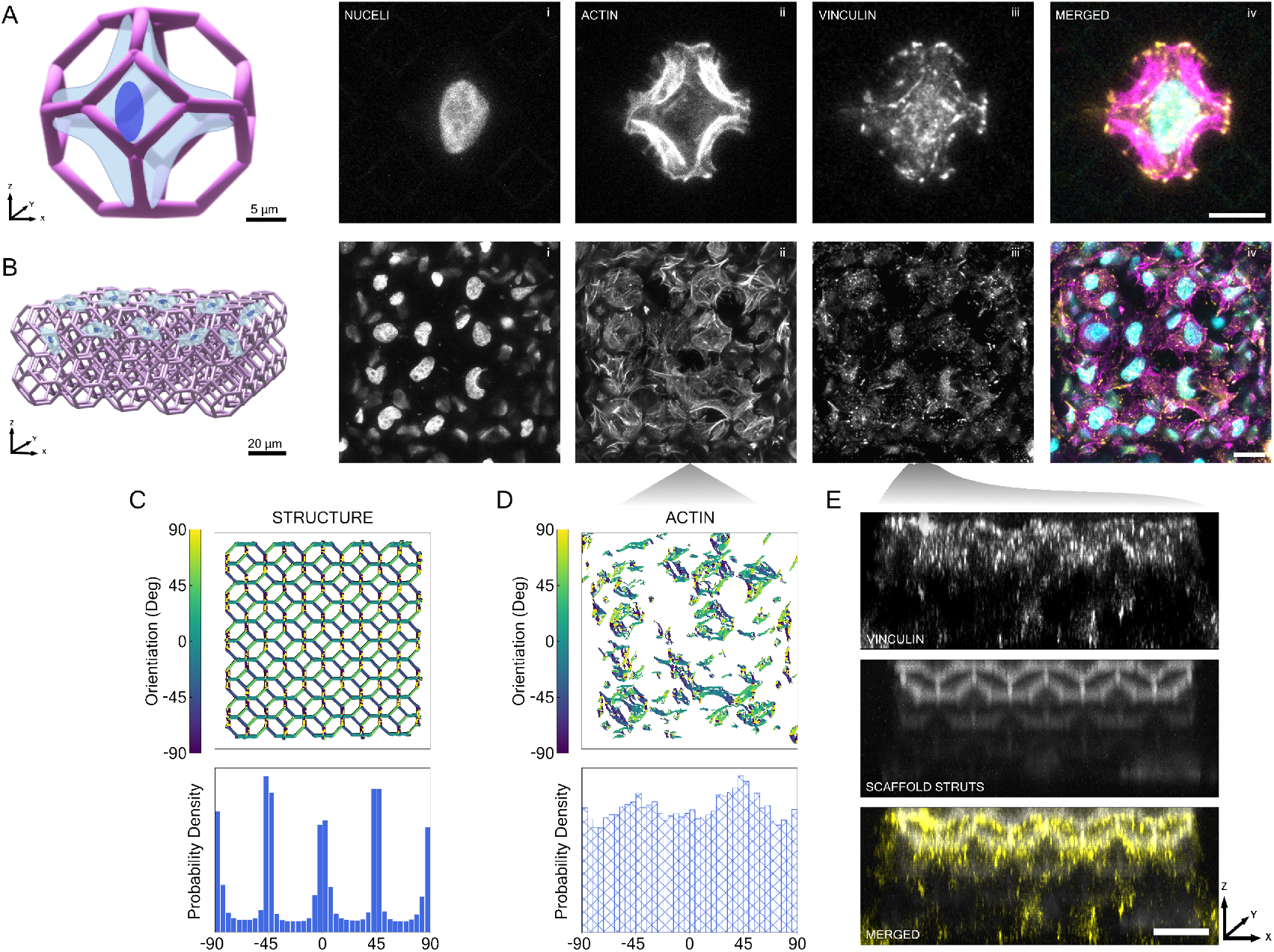
Cell morphology reflects scaffold geometry in absence of loading. A) Representative graphic of isolated cell and its morphology within a structural unit cell. Ai-iv)Confocal maximum intensity projection (MIP) images of a single cell within a unit cell of the nanoarchitecture show spatial adaptation of nuclear morphology (i), actin cytoskeleton arrengement (ii), and focal adhesions distribution (vinculin, iii), with a merged image that shows nuceli (cyan), actin (magenta), and vinculin (yellow) in SAOS-2 cells (iv) (Scale bar = 10 µm). B) Representative graphic of multiple cells cultured on entire nanoarchitecture Bi-iv) Representative MIP images of larger cell populations (up to 37 cells) showing consistent geometric templating (Scale bar = 20 µm). C) Orientation map and orientation distribution of the nanoarchitecture struts from MIP images. D) Segmented actin orientation maps from the population shown in Bii, with the corresponding distribution. E) Projection of the vinculin from Biii showing focal adhesion formed and distributed along the depth of the scaffold geometry, merged image of focal adhesions overlaid struts (Scale bar = 20 µm).

Under low-confluence conditions, an individual cell typically occupied a single unit-cell of the scaffold. DAPI fluorescence staining revealed that nuclei adopted an elliptical shape and were positioned within the voids of the unit cells. Phalloidin staining showed well-defined cortical actin stress fibers aligned along the directions of the struts (Fig. 2A). We also observed prominent focal adhesions at the termini of stress fibers, anchored to the unit cell geometry. At higher confluence, populations of up to 37 cells occupied the 50-unit-cell scaffold (Fig. 2B). Many nuclei retained elliptical shapes, while others showed concavity, likely reflecting local interactions with curved struts. Histograms of nuclear projected area, measured from maximum intensity projection (MIP) images, indicated broader and more uniform distributions compared to cells on flat 2D substrates (Fig. S7). These results suggest that cell and nuclear morphologies adapt to the 3D scaffold geometry, even in dense cultures.

To further quantify how the cytoskeleton conforms to the nanoarchitecture, we measured f-actin orientation and compared it to strut alignment^26^. Figure 2C,D shows the orientation distribution of the scaffold struts, with clear peaks at 45° intervals. The actin filament orientation distribution (Fig. 2D, bottom panel) exhibited similar peaks, indicating alignment with the scaffold geometry. Orientation maps (Fig. 2E) visually confirmed this alignment, while reconstructed cross-sections (Fig. 2F) showed that vinculin-rich focal adhesions were spatially correlated with strut locations. Together, these results demonstrate that the nanoarchitecture guides cytoskeletal alignment and focal adhesion localization through geometric confinement^12^, even in the absence of applied solid or fluid stress.

### Dynamic loading disrupts cytoskeletal symmetry and focal adhesion maturation

To understand how cyclic mechanical loading alters cytoskeletal organization, we fabricated four nanoarchitected scaffolds on a single silicon wafer (Fig. 3A), seeded with SAOS-2 cells, and subjected to compressive strains at varying frequencies. Isolated cells in the individual scaffolds were subsequently imaged; autofluorescence from residual polymer provided a consistent structural reference across all loading conditions. Figure. 3B shows that under static, stress free conditions (0 Hz), actin filaments aligned with the scaffold geometry, recapitulating the symmetry observed in Fig. 2A. With increasing loading frequency (0.1–1 Hz), we observed a breakdown of this symmetry. Confocal images revealed progressively disorganized actin structures, with reduced alignment to scaffold struts (Fig. 3C,D). Orientation histograms confirmed a loss of characteristic *±*45° peaks, indicating a disruption of the lattice-templated actin architecture (Fig. 3E).

**Figure 3:**
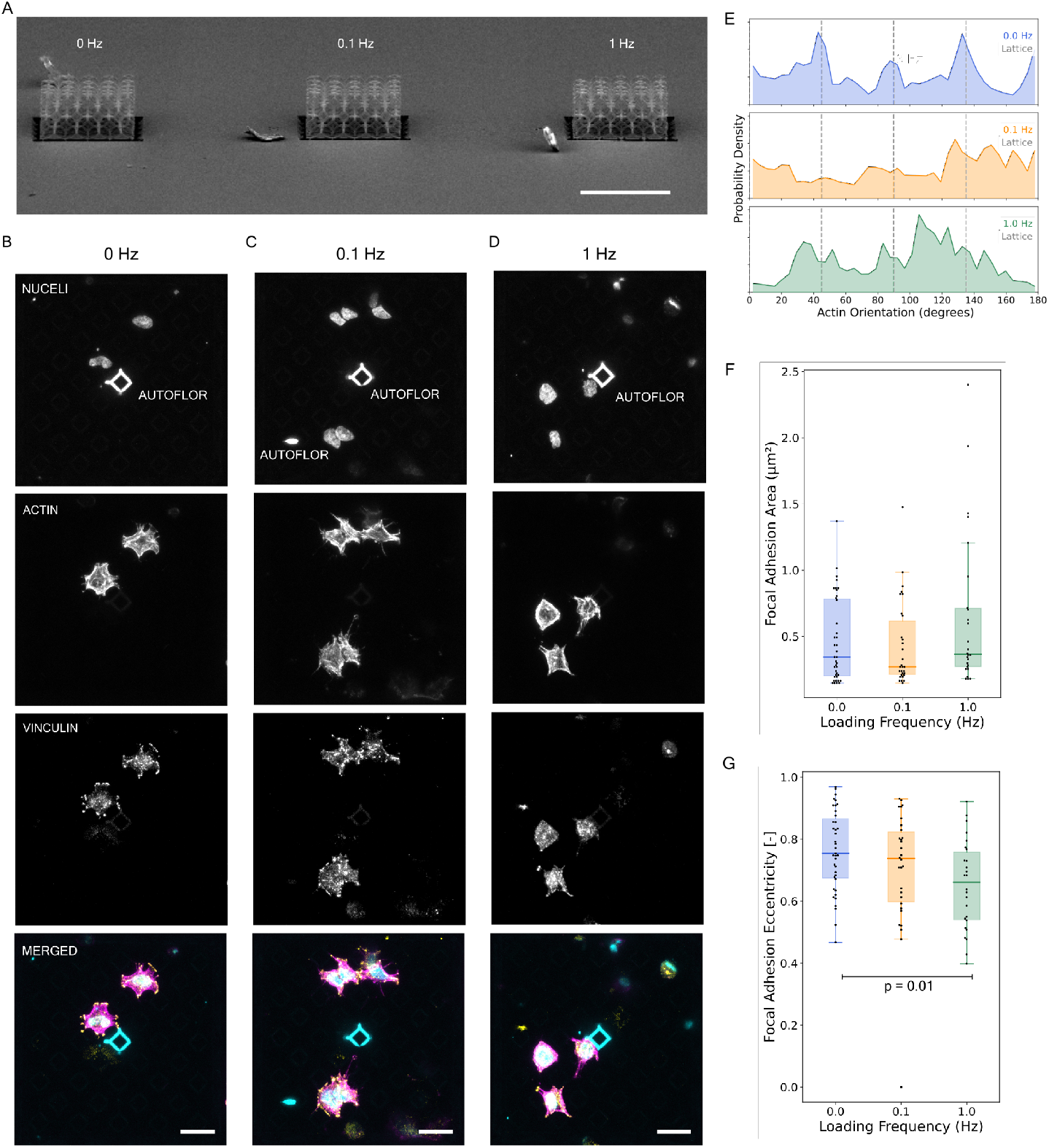
Dynamic loading disrupts cytoskeletal symmetry. A) SEM image of three TiO_2_ scaffolds on a silicone wafer for varying frequencies of dynamic loading (0, 0.1, 1 Hz; Scale bar = 100 µm). (B) In control conditions, actin (magenta) and vinculin (yellow) of isolated cells reflect symmetries of underlying scaffold geometry, visible by residual polymer autoflorescence. Loading of cell-laden scaffolds at 0.1 Hz C) and 1 Hz D) results in disruption of of symmetry and qualitative reduction of focal adhesions (Scale bar = 20 µm). E) Orientation histograms of actin filaments (shaded) show loss of structural lattice peaks (*±*45°, dashed lines) with increased loading frequency. F,G) Areas and eccentricities of individual focal adhesions, measured via vinculin, as a function of loading Eccentricity decreases with increasing loading frequency, consistent with less mature focal adhesion.

To assess whether focal adhesions exhibited similar remodeling, we segmented vinculin-stained adhesions and quantified both area and eccentricity. While adhesion area remained relatively constant, eccentricity declined significantly with increasing frequency (Fig. 3F,G), suggesting a shift from elongated, stable adhesions toward smaller, more punctate forms. Because elongation and eccentricity are hallmarks of adhesion maturation^27^, these results point to a lack of adhesion maturation process under cyclic loading. Collectively, these findings suggest that dynamic biphasic loading alters both cytoskeletal and adhesive organization, even at low frequencies. The progressive loss of alignment and adhesion eccentricity suggests that fluid-solid coupling may serve as a mechanical signal that primes cells for reorganization, potentially toward a migratory or remodeling phenotype.

### Cells sense and respond to subtle increases in solid–fluid coupling

Cells embedded in 3D extracellular environments are known to integrate mechanical inputs from both the solid matrix and the interstitial fluid. To quantify how dynamic biphasic stimuli affect cytoskeletal organization, we computed a dimensionless frequency 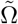 described in Eq. 1

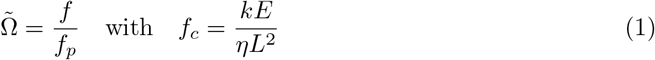

where *f* is the loading frequency, and 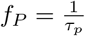 where *τ*_*p*_ is the poroelastic relaxation time that reflects the natural drainage rate of the scaffold^28,29^. It depends on the permeability *k*, effective modulus *E*, fluid viscosity *η*, and a characteristic structural length scale *L*. The parameter 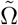 provides a measure of how rapidly the scaffold is being deformed relative to the rate at which fluid can redistribute through it. Small values where 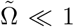 indicate the “drained” regime in which fluid flow is relatively unrestricted by the porous structure; as 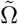 approaches 1, fluid stress and solid deformation become more coupled. We place our results in this poroelasticity context by calculating 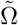 using *K* = 10^−13^ cm^−2^, *η* = 10^−3^ Pa · s, *E* ∈ [2.4 7] MPa and *L* ≈ 25 µm.

In Fig. 4A,B we relate the calculated 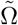 to average actin alignment to the scaffold^11^, defined as the order parameter ⟨*S*_45_⟩ (See Methods). We observe that across all scaffold conditions spanning a range of stiffnesses (2.4–7 MPa) and frequencies (0–3 Hz), there is monotonic decline in actin alignment as 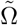 increased from 10^−6^ to 10^−5^ (Fig. 4A,B). Importantly, this disruption of scaffold-guided cytoskeletal alignment occurred entirely within the “drained” regime where 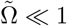, suggesting that cells are highly sensitive to even small departures from equilibrium fluid conditions. This sensitivity implies that cells may perceive subtle increases in fluid shear or pressure as cues for remodeling their cytoskeleton.

**Figure 4:**
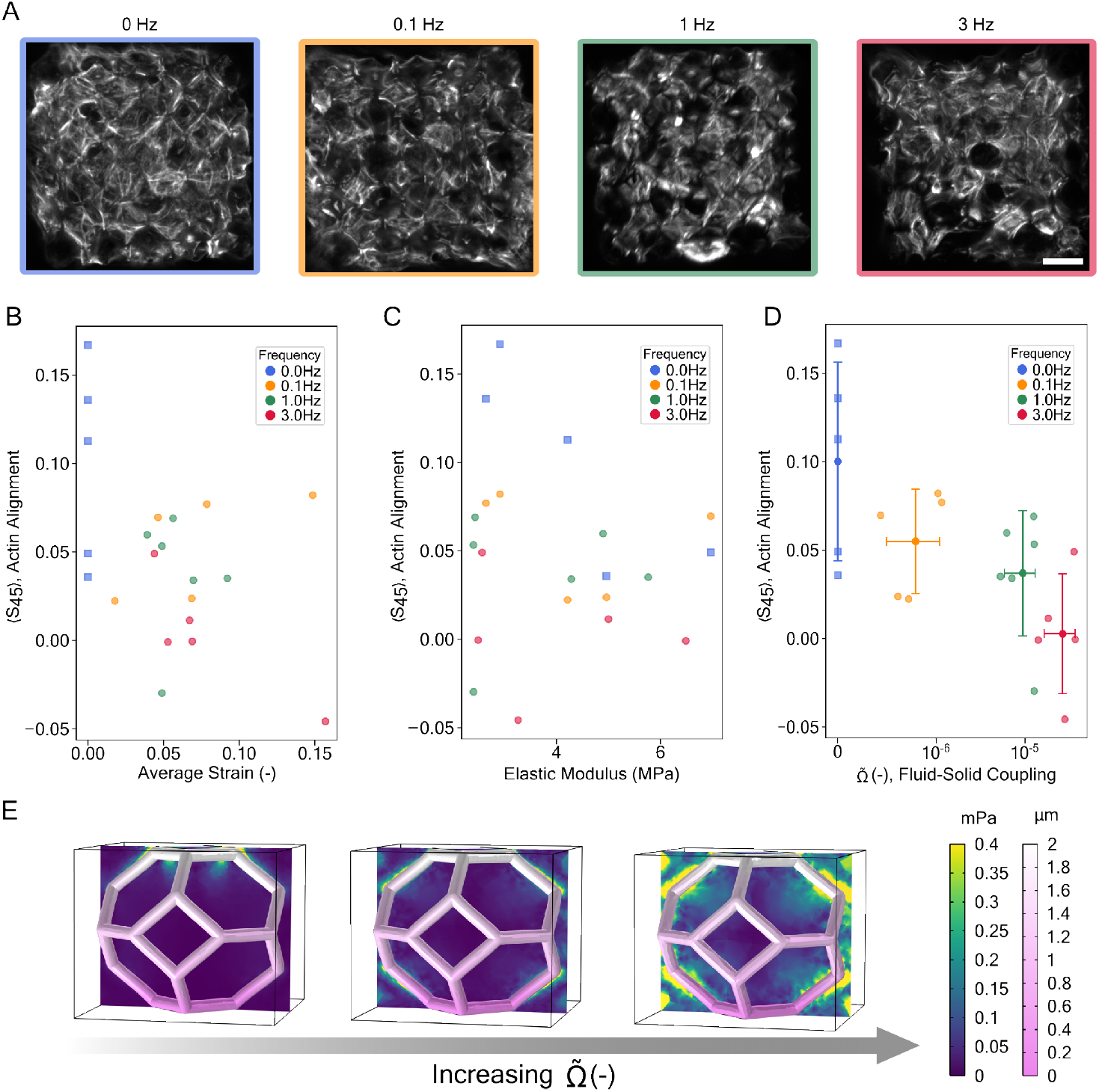
Cystoskeletal order parameter in dynamic poroelastic loading. A) Decrease in actin alignment with increasing of loading frequency. B) No correlation between order parameter ⟨*S*_45_⟩ and average strain as lattice cyclic compression displacement increases. C) No correlation between order parameter ⟨*S*_45_⟩ and elastic modulus as compression frequency increases. D) Order parameter ⟨*S*_45_⟩ decreases with increase coupling parameter 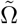, indicating greater cell sensing of solid-fluid interactions in poroelastic extracellular environment. E) Finite Element simulations show increase in coupling 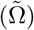 correlates with high fluid shear stress near concentration near struts where focal adhesions form.

Simulations confirmed that increasing the loading frequency elevates localized fluid shear stress, with values approaching 0.4 mPa at 3 Hz (Fig. 4C). These stresses are concentrated near lattice struts, precisely where focal adhesions tend to form, and may modulate intracellular tension or disrupt cytoskeletal anchoring. Along with the alignment data, our computational results indicate that cells gradually remodel their cytoskeleton in response to even modest increases in solid–fluid coupling.

## Discussion

In this study, we used architected, biphasic scaffolds to dissect how cells respond to combined matrix and fluid mechanical cues in a 3D environment. Our results reveal that cells seeded on static scaffolds adopt cytoskeletal and nuclear morphologies that mirror the underlying geometry, with actin fibers aligning to lattice struts and nuclei suspended or deformed depending on local curvature (Fig. 2). This templating behavior occurs in the absence of external loading, indicating that geometry alone is suffcient to organize subcellular structure, likely through spatial confinement and adhesion patterning^30–32^. Focal adhesions are localized to scaffold struts, providing anchorage points that define cortical actin tension and guide alignment. These results highlight the instructive role of architecture in shaping cell behavior in 3D.

By adding cyclic loading, we observed frequency-dependent disorganization of the actin cytoskeleton and loss of focal adhesion eccentricity, with changes emerging even at 0.1 Hz (Fig. 3). These shifts reflect a disruption of the scaffold-guided symmetry established under static conditions. While total focal adhesion area remained constant, their reduced eccentricity under dynamic loading suggests a transition from mature, elongated adhesions to more punctate, transient forms, consistent with a dematuration process^27^. Because mature adhesions stabilize stress fibers and resist traction, their disassembly is a prerequisite for motility, suggesting that dynamic loading may prime cells for migration even in the absence of directional cues. These findings are consistent with prior work showing that focal adhesion maturation is governed not only by the magnitude of applied tension but also by the spatial and temporal context in which it is delivered. It has been demonstrated that myosin-generated tension alone is insuffcient to drive elongation unless stress fibers provide a pre-aligned actin template to guide adhesion assembly^33^. Similarly, force-dependent adhesion growth requires sustained loading over specific timescales to trigger reinforcement, with insuffcient duration failing to support maturation^34^. Our observation that focal adhesion eccentricity declines under higher-frequency loading suggests that cyclic mechanical inputs at short timescales disrupt these stabilizing templates, preventing the assembly of mature adhesions and thus biasing cells toward disorganized, motile states. These dynamics may explain why cytoskeletal symmetry degrades under fluid-constrained loading even without large deformation amplitudes.

To mechanistically unify observations across stiffness and frequency regimes, we introduced nondimensional 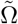, which quantifies the relative rate of loading to fluid redistribution. We found that actin alignment declines monotonically as 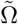 increases, even though all values remained well below unity (Fig. 4). Prior studies have modeled frequency-dependent reorientation using exponential dynamics or stress fiber turnover rates modulated by amplitude and substrate mechanics^11,35^. In contrast, 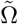 captures the interplay between matrix strain rate and fluid drainage, enabling comparison across dynamic loading regimes. This surprising sensitivity suggests that cells respond to small departures from equilibrium drainage, possibly via local shear stresses or pressure gradients generated at cell–scaffold interfaces. Simulations revealed peak shear stresses on the order of 0.4 mPa at 3 Hz (Fig. 4C), substantially lower than those known to drive remodeling in 2D monolayers^36^, and implicating solid–fluid coupling as a biologically meaningful input in 3D mechanosensing.

This correlation across diverse loading conditions suggests that poroelastic coupling could provide a broader context for interpreting cytoskeletal responses to dynamic mechanical environments. While poroelasticity has been studied extensively in tissue-scale biomechanics^1,2,37^, few studies have linked it to subcellular organization^28^. Our findings show that macroscopic mechanical inputs can be transformed into biologically relevant signals through cell-scale solid–fluid interactions, particularly in architected matrices where geometry governs fluid redistribution and mechanical anisotropy.

Importantly, these insights arise from an experimental system that allows direct control over scaffold architecture, matrix stiffness, and loading parameters.In tissues such as bone, cartilage, and developing vasculature, cells experience cyclic loading within confined porous environments, conditions recapitulated in our system^6,9,38^. The ability to resolve how cells transition from symmetry to disorganization, and from stable adhesion to potential migration, opens the door to deeper investigations of mechanotransductive signaling in physiological-like settings.

### Outlook and Applications in 3D Mechanobiology

This work establishes a tunable platform to investigate how cells integrate solid matrix deformation and interstitial fluid flow within a 3D nanoarchitected environment. By combining in situ imaging, focal adhesion quantification, and poroelastic modeling, we demonstrate that small increases in a dimensionless solid–fluid coupling parameter, 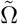, are suffcient to disrupt actin symmetry templated by scaffold geometry and to reduce focal adhesion eccentricity, a hallmark of maturation. That this loss of cytoskeletal order occurs while 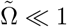 underscores the sensitivity of cells to subtle departures from equilibrium drainage conditions.

Because focal adhesions form near scaffold struts, precisely where simulated shear stresses concentrate, our results suggest that dynamic loading introduces local mechanical gradients that interfere with cytoskeletal stability. This biphasic mechanical context may help explain how cells initiate cytoskeletal remodeling or migration in response to dynamic loaded environments like bone^39^, cartilage^40^, or developing morphogenetic structures^41^.

Future efforts would further explore how spatiotemporal anisotropy, such as directional gradients in fluid–solid coupling, could control phenotypic transitions or alignment states in 3D^42^. These approaches could build on recent insights into anisotropic mechanosensing^43^, where directional stress reinforcement stabilized cytoskeletal symmetry and activation. In particular, the ability to tune scaffold stiffness, porosity, and architecture while delivering defined biphasic loads opens new opportunities to decouple competing mechanical stimuli and identify critical thresholds for mechanosensitive signaling in 3D.

## Materials and Methods

### Scaffold fabrication

The scaffold was printed on a silicon wafer using two-photon lithography technology with the Nanoscribe Photonics Professional GT. The 1 cm silicon wafer featured an inner 7 cm × 7 cm square well that etched to a depth of approximately 300 µmto hold cell culture media. Four scaffolds, each 125 µm× 125 µm× 50 µm, were printed in a row within this etched area. The polymer scaffolds were coated with a ∼50 nm layer of TiO_2_ using the Cambridge Nanotech S200 ALD. The edges of the TiO_2_-coated scaffolds were then cut to expose the polymer with the Tescan S8000X Plasma Focused Ion Beam. Following approximately 32 hours of oxygen plasma etching, titania scaffolds with hollow struts were obtained.

### COMSOL simulation model

To simulate the compression-induced fluid shear stress with finite element method in COMSOL, we prescribed the top surface of a unit cell with a velocity that corresponds a 1 Hz compression cycle with maximum displacement of 2 *upmu*m, while the bottom surface remained fixed. The corresponded fluid stress field, represented by the cubic frame in Fig. 1D was calculated with the following boundary conditions: the top surface followed the solid displacement, the bottom surface was fixed, and all other surfaces were assigned Dirichlet boundary conditions for moving mesh and set to be fluid outlet. Initially, the fluid within the system was at rest, exhibiting no movement or flow. As compression was applied, the fluid (water) was displaced and flowed outward through the four side faces of the cubic domain. The fluid-solid interaction surfaces adhered to no-slip boundary conditions.

### Cell culturing

The SAOS-2 cell line from ATCC was used in all in-vitro experiments. Cells were cultured in 75 cm^2^ and 175 cm^2^ tissue culture flasks. The cell culture media comprised Dulbecco’s Modified Eagle’s Medium (DMEM), 10% fetal bovine serum (FBS), and 1% penicillin-streptomycin (10,000 U/mL). The media was filtered through a 1000 mL filter unit with a 0.2 *mum* aPES membrane and replaced every 2-3 days. Cells were passaged upon reaching 80% or more confluence using Accutase Cell Detachment Solution, then either seeded into new tissue culture flasks or onto the scaffolds. Before compression tests, cells were seeded onto the TiO_2_ scaffolds at a density of cultured for at least three days in growth media, with a media change on the second day. Compression experiments were conducted on day 3 with cells attached to the scaffolds.

### Mechanical stimulation

The compression tests were conducted using the KLA Nanoindenter equipped with a 150 µmflat-punch diamond tip. Prior to compressing the scaffold, the experimental station and tool were sanitized with 70% ethanol. Cyclic compression tests were applied with a load range of 0.5 mN to 1 mN and a frequency range of 0.1 Hz to 3 Hz for 80 cycles. Additional loading at 1.5 mN was performed, which resulted in complete or near-complete cell detachment in all applied frequency groups (Fig. S8). A dynamic control method was used during the tests to ensure proper contact without compromising the resolution of stress and strain measurements.

### Immunofluorescent staining and imaging

After the compression tests, samples were washed twice with phosphate-buffered saline (PBS) and then fixed with 4% paraformaldehyde for 10-15 minutes. The paraformaldehyde was subsequently washed away with two PBS washes. Following the PBS washing, samples were blocked with 1% BSA in PBS for 30 minutes. After blocking, an anti-vinculin primary antibody, diluted in blocking buffer, was applied to the cells and incubated overnight at 4°C. The samples were then washed three times with PBS before being treated with a vinculin-secondary antibody for 90 minutes at room temperature. After additional PBS washes, cytoskeletal actin and cell nucleus staining were performed by incubating with conjugated phalloidin-555 (SantaCruz) for 90 minutes and DAPI 488 for 10 minutes at room temperature, respectively. Following a final PBS wash, the sample substrates were mounted onto glass slides using Fluoromount-G mounting media. Cell imaging was conducted using a Zeiss LSM 710 confocal microscope. Using a 20X objective, z-stack image scans were collected with an interval distance of 0.15 µm.

### Quantification of actin alignment to scaffold geometry

To quantify how well actin filaments aligned with the underlying scaffold geometry, we computed an orientation order parameter ⟨*S*_45_⟩ = ∑*h*(*θ*) cos(8*θ*), where *h*(*θ*) is the normalized histogram of actin fiber orientations and *θ* is the angle in radians. The function cos(8*θ*) captures alignment to the scaffold’s octagonal symmetry, with preferred orientations at 0°, *±*45°, and *±*90°. An ⟨*S*_45_⟩ value of 1 indicates perfect alignment to these directions, 0 reflects a random orientation distribution, and -1 indicates alignment at angles orthogonal to the scaffold symmetry.

## Acknowledgments

This work is supported by O.A.T’s NSF CAREER Award CMMI - 2339836. This work was supported in part by the Center for Engineering MechanoBiology, NSF Science and Technology Center CMMI: 15-48571 We sincerely thank the Singh Center for Nanotechnology and the Beckman Center for Cryo-Electron Microscopy at the University of Pennsylvania for their invaluable assistance and access to their tools and facilities. This work is supported by the Cell & Developmental Biology (CDB) Microscopy Core (RRID SCR022373) at the University of Pennsylvania. We thank Alessandro Maggi for helpful discussions. We sincerely thank Andrea Stout and Jasmine Zhao for their time in troubleshooting microscopy. The work was supported (in part) by a seed grant from the Center for Precision Engineering for Health (CPE4H) from Penn Engineering, University of Pennsylvania.

## Author contributions

KC, ABC, MLP, LC, EB, TW and RT performed the experiments. KC performed the simulations. KC, ABC, MLP, TW and OAT analyzed the data and wrote the paper. All authors discussed the results and revised edited manuscript. OAT supervised the study.

