## Supporting Information for "Biphasic Mechanical Loading Disrupts Cytoskeletal Symmetry in 3D Architected Scaffolds"

Supporting Information

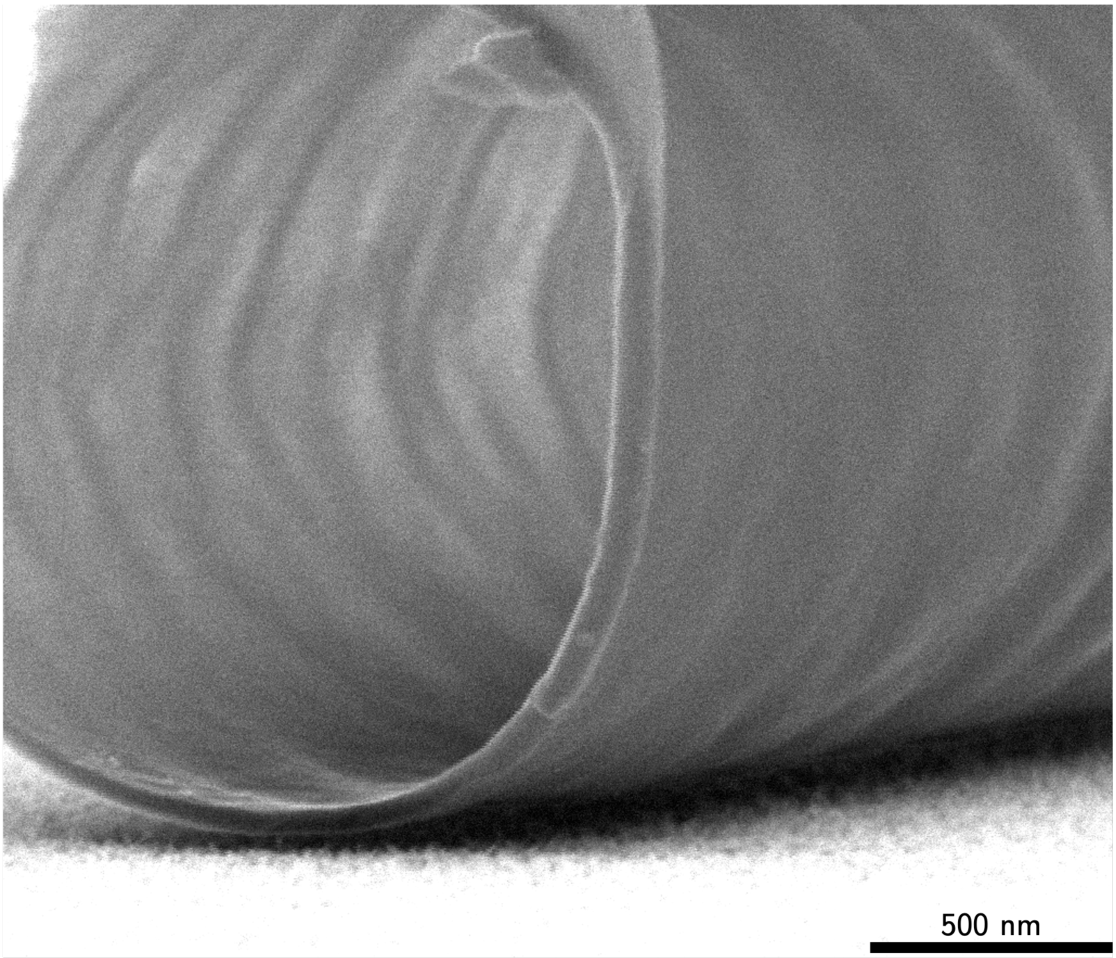


**Figure S1:** SEM imaging showing a single hollow strut with a $TiO_2$ wall after the removal of the polymer inside. Wall thickness is approximately 50nm.

### COMSOL simulation of parallel flow around the nanoarchitecture

To simulate the flow field around the nanoacrchitecture, a laminar parallel flow model was applied in the volume of the cube with the structure in the center. The water flows into the cube with a constant velocity of 10 $\mu m/s$ from plane X=0. All other faces are set to be non-slip. The streamline shows some curvature near the struts, as the µPIV test shows in Fig. 1C, while the flow velocity slows down inside the structure and results in a lower outlet velocity. Note that boundary layers were formed at the gap between the prescribed water boundary and the structure edge, but no obvious boundary layers were observed in the µPIV test due to a much larger channel design.

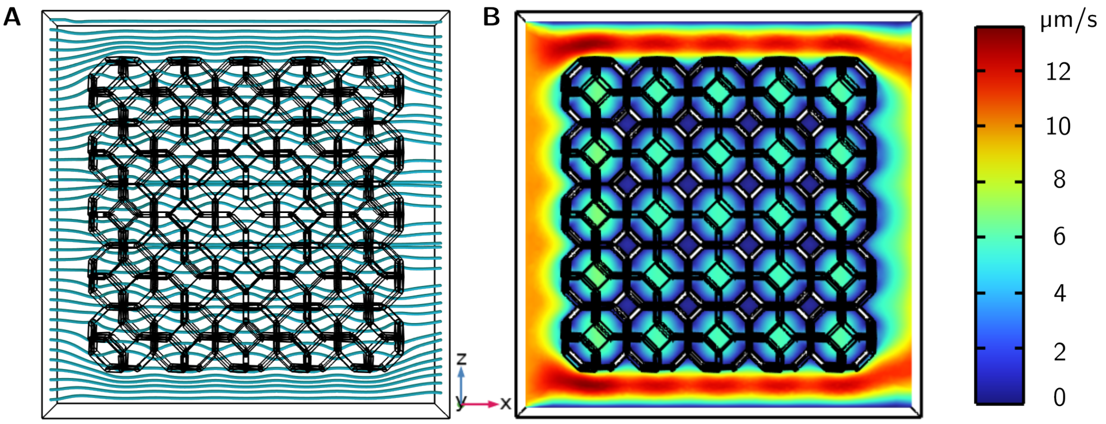


**Figure S2:** A. Streamline and B. Velocity Field around the lattice with parallel flow towards positive x direction

### Theoretical squeezing flow model

Compression of the high porosity scaffold is simplified as a two-dimensional squeezing flow model between two infinitely long parallel plates, whose scheme is shown in Fig.S3. The loading process on the stiff scaffold immersed in culture media is simulated by the approaching top and bottom plates relative to the center axis, with a velocity of 2.5 $\mu m/s$, chosen reasonably for the following calculation based on previous loading tests (need more details or reference based on the contents). The fluid between the parallel plates is squeezed out as two plates moving. Correspondingly, the governing equations of the fluid in between are written as

$$\frac{\partial u}{\partial x} + \frac{\partial v}{\partial y} = 0 \tag{2}$$

$$\frac{\partial u}{\partial t} + u\frac{\partial u}{\partial x} + v\frac{\partial u}{\partial y} = -\frac{1}{\rho}\frac{\partial P}{\partial x} + \nu(\frac{\partial^2 u}{\partial x^2} + \frac{\partial^2 u}{\partial y^2}) \tag{3}$$

$$\frac{\partial v}{\partial t} + u\frac{\partial v}{\partial x} + v\frac{\partial v}{\partial y} = -\frac{1}{\rho}\frac{\partial P}{\partial y} + \nu(\frac{\partial^2 v}{\partial x^2} + \frac{\partial^2 v}{\partial y^2}) \tag{4}$$

where $u$ is the fluid velocity component along $x$ direction, $v$ is the fluid velocity component along $y$ direction, $t$ is time, $\rho$ is fluid density, $\nu$ is fluid kinematic viscosity, $P$ is pressure. Note that here we assume the fluid is viscous and incompressible. The boundary conditions are

$$y = a(t) : u(x, y, t) = 0; v(x, y, t) = V_w(t) \tag{5}$$

$$y = 0 : \frac{\partial u(x, y, t)}{\partial y} = 0; v(x, y, t) = 0 \tag{6}$$

where $V_w(t) = da/dt$ is the upper plate velocity. To obtain a self-similar solution[44], it is assumed that

$$u(x, y, t) = \frac{C - x}{a(t)} V_w(t) f^{'}(\eta) \tag{7}$$

$$v(x, y, t) = V_w(t) f(\eta) \tag{8}$$

where $\eta = y/a(t)$ and Eq. 5 and 6 satisfy the continuity equation, cf. Eq. 1. Here C is a constant related to inlet condition of the channel and is set to be 0 in the current case since the scaffold is submerged in a static well at t=0. Substituting Eq. 5 and 6 into Eqs. 1-3, we have

$$R(ff^{'''} - f^{'}f^{''} - \eta f^{'''} - 3f^{''}) = f^{iv} \tag{9}$$

where $R = a(t)V_w(t)/\nu$ is the Reynolds number. The boundary conditions become

$$\eta = 0 : f(0) = 0; f^{''}(0) = 0 \tag{10}$$

$$\eta = 1 : f(1) = 1; f^{'}(1) = 0 \tag{11}$$

By employing Runge-Kutta 4th order method, Eqs. 7-9 can be solved, while the missing two initial conditions, $f^{'}(0)$ and $f^{'''}(0)$, are found by the shooting method. The solutions of $f$, $f^{'}$, $f^{''}$ and $f^{'''}$ at the time moment when $a(t) = 40\mu m$, are shown in Fig. S4. Note that here we took $a(t = 0) = 50\mu m$. The resulted Reynolds number is in the order of $10^{-7}$, which indicates laminar flow. As is seen, the $x$-component velocity $u$ at certain $x$, represented by $f^{'}(\eta)$, decreases hyperbolically from the maximum at the center axis to 0 at the top plate. The $y$-component velocity $v$, represented by $f(\eta)$, increases linearly from 0 at the center axis to the maximum at

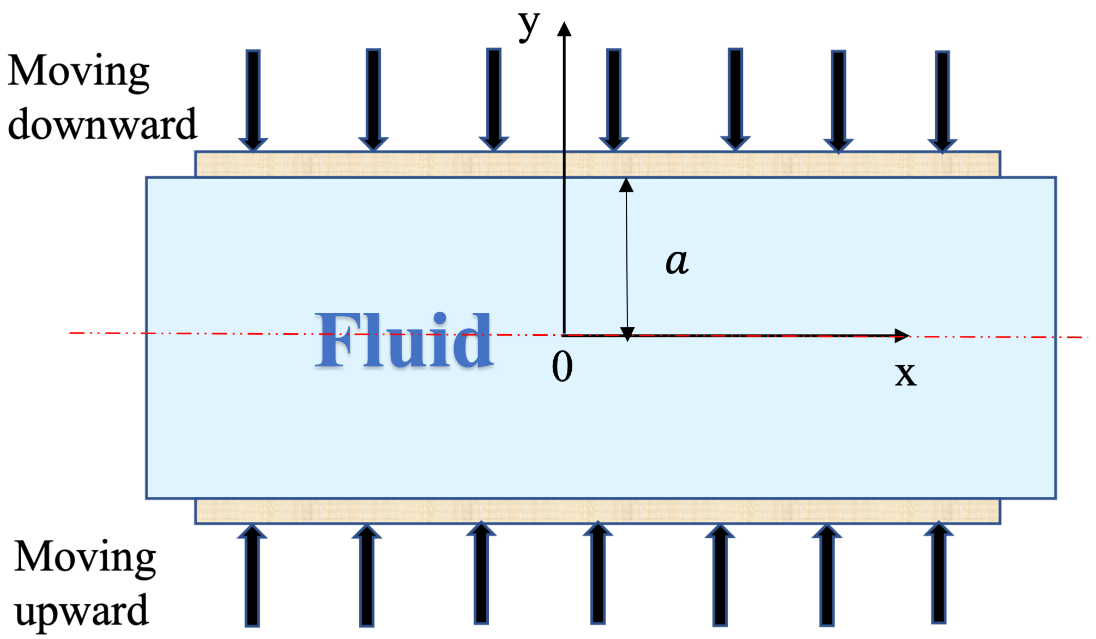


**Figure S3:** Scheme of squeezing flow between two parallel plates.

the top plate. Same tendency can also be seen in Fig. S5, where the horizontal velocity $u$ and vertical velocity $v$ in a $50\mu m$ by $50\mu m$ area in the first quadrant of the $Oxy$ coordinate system are plotted. Accordingly, the shear stress, shown in Fig. S6b, is calculated by the below equation

$$\tau = \mu \frac{\partial u}{\partial y} \tag{12}$$

where $\mu$ is the fluid dynamic viscosity. The maximum shear stress is at the upper corner where the horizontal velocity gradient is the largest, and the value is around 0.2 $mPa$ with the plate moving velocity of 2.5 $\mu m/s$. This reveals different forces sensed by the seeded cells at different locations during the loading process. In addition, shear stresses induced by the plate moving with velocity of 0.25 $\mu m/s$ and 7.5 $\mu m/s$ are also calculated as shown in Fig. S6a and S6c. It is seen that with faster plate moving velocity, shear stress at the same spot increases correspondingly, which may indicate different cell responses under different loading conditions.

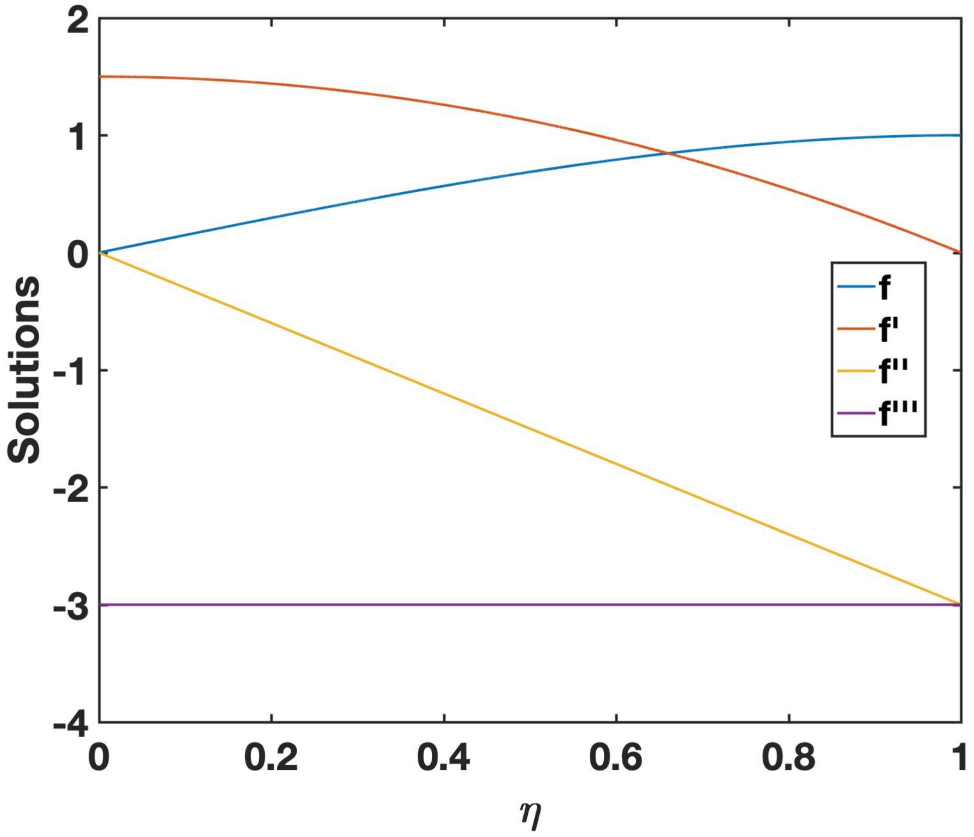


**Figure S4:** Dimensionless solutions.

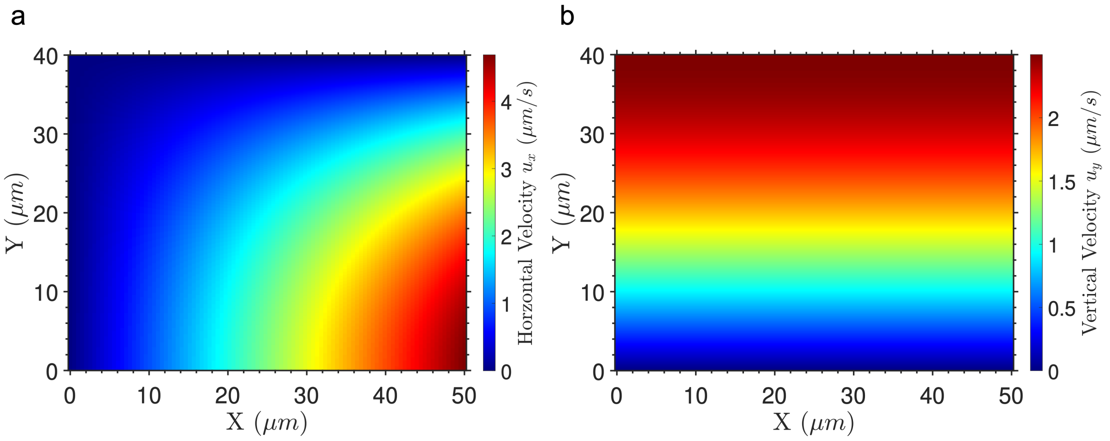


**Figure S5:** Velocity components along (a) $x$ direction and (b) $y$ direction in the area of $0 \leq x \leq 50\mu m$, $0 \leq y \leq 50\mu m$.

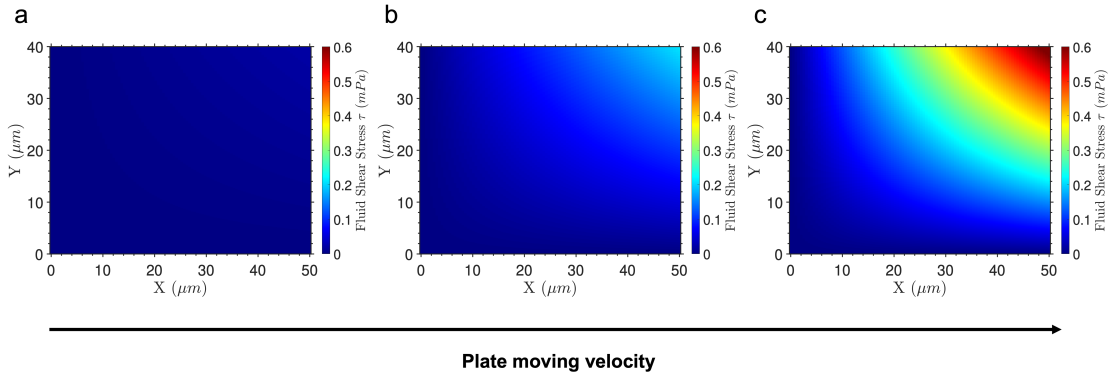


**Figure S6:** Shear stress field in the area of Fig. S5 with plate moving velocity of (a) 0.25 $\mu m/s$, (b) 2.5 $\mu m/s$, and (c) 7.5 $\mu m/s$.

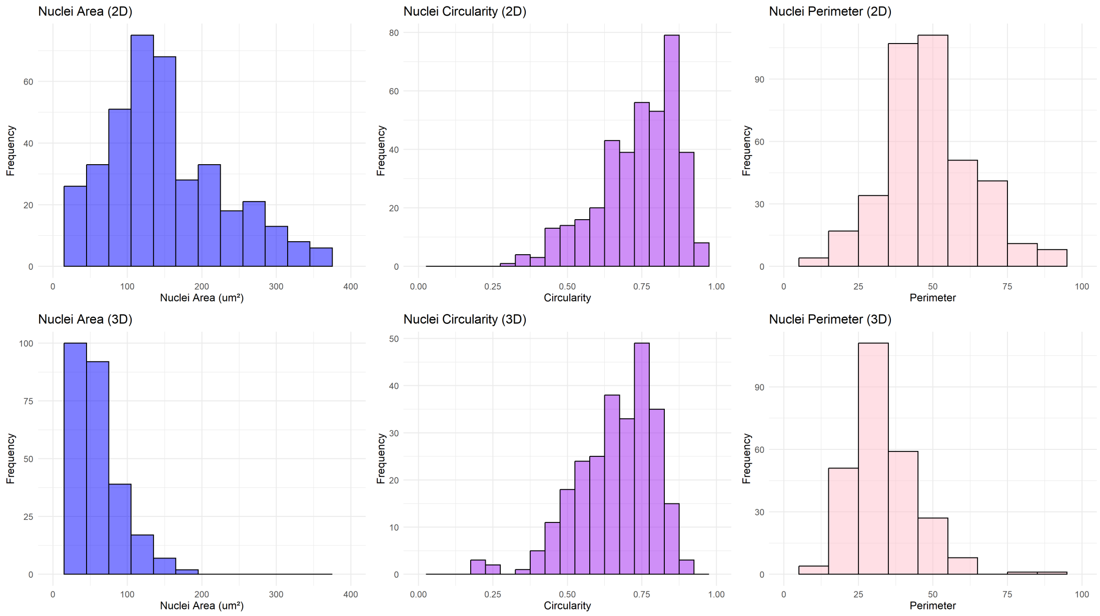


**Figure S7:** Histograms of nuclear area, circularity, and perimeter for SAOS-2 cells cultured on 2D substrates (top row) and within 3D TiO2 lattices (bottom row). Nuclear morphology was quantified from maximum intensity projections (MIP) of the DAPI channel from zStacks using a custom ImageJ macro. Images were converted to 8-bit, Gaussian blurred (sigma = 0.5 for 2D, sigma = 2 for 3D), thresholded using the "Default" method, and processed with watershed segmentation. The "Analyze Particles" function was run with a custom size (30–450 um2 and circularity = 0.00–1.00) to help exclude noise. Nuclear circularity was then computed with the area and perimeter values in Excel using the formula: $Circularity = 4\pi\frac{Area}{Perimeter^2}$

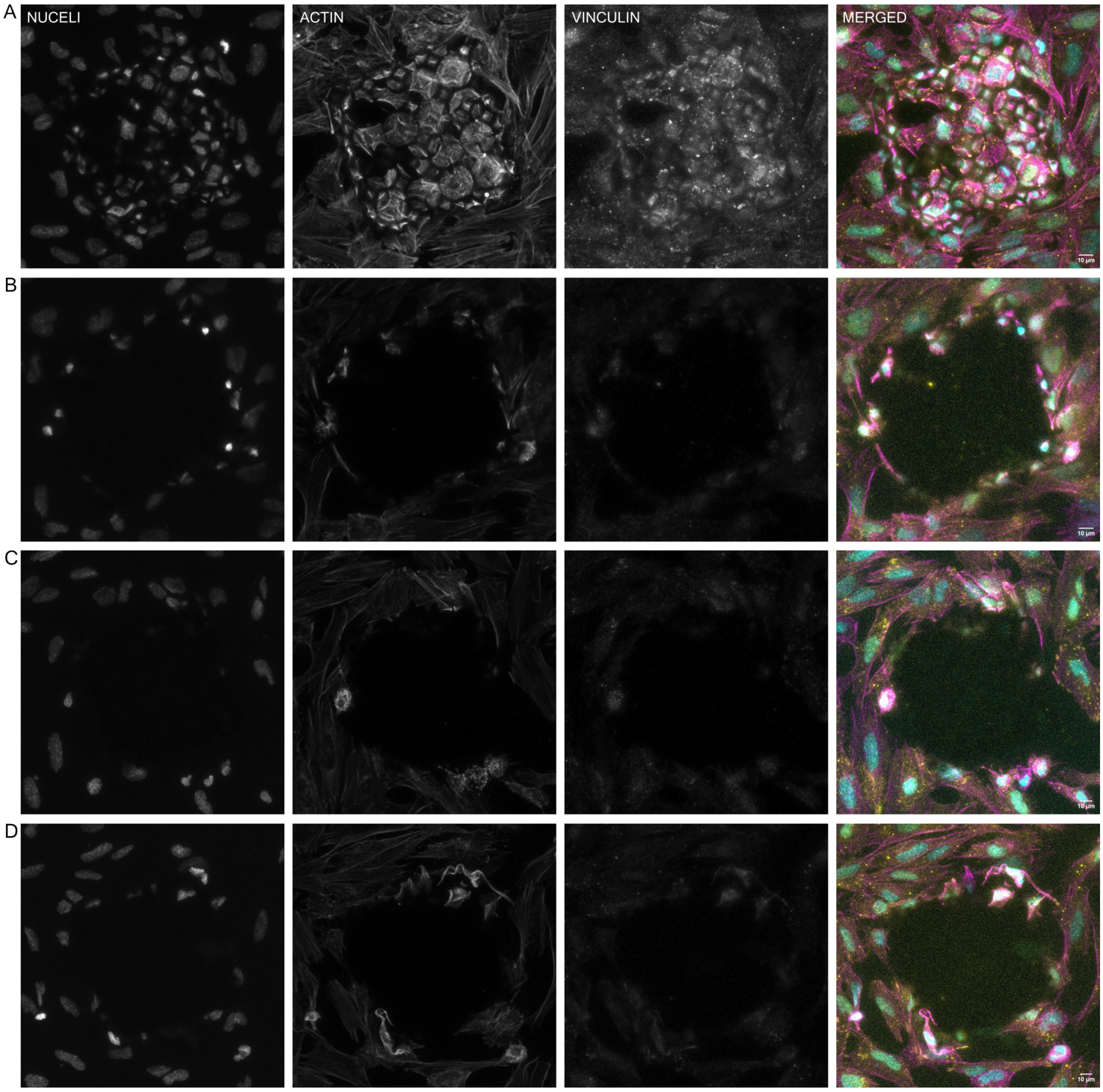


**Figure S8:** Confocal MIP images showing nuclei, actin, vinculin, and merged channels for A. 0 Hz, B. 0.1 Hz, C. 1 Hz, and D. 3 Hz experimental groups. Samples were exposed to cyclic loading with a peak force of 1.5 mN. In the 0.1–3 Hz groups (B–D), cells are observed only on the surrounding 2D substrate, with little to no cells remaining on the $TiO_2$ lattices following compressions. Scale Bar = $10\mu m$. (Merged channel legend; cyan = nuclei, magenta = actin, yellow = vinculin)
